# Do mixed-species groups travel as one? An investigation on large African herbivores using animal-borne video collars

**DOI:** 10.1101/2024.04.09.588700

**Authors:** Romain Dejeante, Marion Valeix, Simon Chamaillé-Jammes

**Affiliations:** CEFE, Univ Montpellier, CNRS, EPHE, IRD, Montpellier, France; Mammal Research Institute, Department of Zoology and Entomology, University of Pretoria, Pretoria, South Africa

**Keywords:** animal movement, heterospecifics, movement ecology, mutualism, landscape of fear, predator-prey interactions

## Abstract

Although prey foraging in mixed-species groups benefit from a reduced risk of predation, whether heterospecific groupmates move together in the landscape, and more generally to what extent mixed-species groups remain cohesive over time and space remains unknown. Here, we used GPS collars with video cameras to investigate the movements of plains zebras (*Equus quagga*) in mixed-species groups. Blue wildebeest (*Connochaetes taurinus*), impalas (*Aepyceros melampus*) and giraffes (*Giraffa camelopardalis*) commonly form mixed-species groups with zebras in savanna ecosystems. We found that zebras adjust their movement decisions solely to the presence of giraffes, being more likely to move in zebra-giraffe herds, and this was correlated to a higher cohesion of such groups. Additionally, zebras moving with giraffes spent longer time grazing, suggesting that zebras follow giraffes to forage in their proximity. Our results provide new insights on animal movements in mixed-species groups, contributing to a better consideration of mutualism in movement ecology.

## INTRODUCTION

Interactions with heterospecifics may affect animal movement decisions. An obvious example is how prey respond to an encounter with a predator (Courbin et al. 2016). Although interspecific interactions are much more diverse than predator-prey interactions, ecologists mainly focus on the influence of the negative interspecific interactions (such as predation or competition) on animal movement ecology (Shaw et al. 2021). Therefore, little is known about how positive interspecific interactions, in particular mutualism, may motivate or constraint movement decisions of individuals (Shaw et al. 2021).

Mixed-species groups are well-known examples of positive interactions between prey: by associating with heterospecifics, individuals benefit from an increased chance of detecting the predator early, and a general dilution of the risk of being attacked (Goodale et al. 2019, 2020). Although the benefits of mixed-species groups have received a lot of attention, whether heterospecific groupmates move together in the landscape, and more generally to what extent mixed-species groups remain cohesive over time and space remains commonly unknown (Carlson et al. 2023). Heterospecific groupmates may have different energetic needs or different habitat selectivity that potentially constrain differently their movement decisions and that may therefore constrain the cohesion of mixed-species groups over time. For example, in savanna ecosystems, plains zebras (*Equus quagga*) and blue wildebeest (*Connochaetes taurinus*) are often observed foraging together, with reduced vigilance compared to when foraging alone (Schmitt et al. 2014). Although they sometimes migrate together (Anderson et al. 2024), their fine-scale movements are differently constrained by habitat selectivity (zebras being more generalist grazers than wildebeest), with known consequences on their spatial responses to the short term risk of predation (Martin and Owen-Smith 2016), but unknown consequences on the cohesion of the mixed-species groups they form.

The knowledge gap in our understanding of the link between positive interspecific interactions and animal movements (Shaw et al. 2021) likely comes from the difficulty to simultaneously, and over time, track the movements of animals and the context in which they occur, in particular the presence or absence of other species. Although large deployment of GPS collars can sometimes allow the simultaneous investigation of spatial and social behaviours (Wilmers et al. 2015), many limitations remain: collars generally do not provide information on the fine-scale behaviour of the tracked individuals (e.g. foraging vs. vigilance) nor on the presence of untracked individuals. Therefore, complementary tools are required. In particular, animal-borne video cameras could fill this methodological gap as they provide data to understand the social context and interactions of tracked animals (Yoda et al. 2011; Bombara et al. 2017; Kaczensky et al. 2019).

Here, we used GPS collars equipped with video cameras to investigate whether plains zebras, who are regularly observed in mixed-species groups in African savannas (Schmitt et al. 2014, 2016; Stears et al. 2020) (see Figure 1 for an illustration of a mixed-species group seen through the eyes of a collared zebra) do travel with heterospecifics in the landscape.

**Figure 1.**
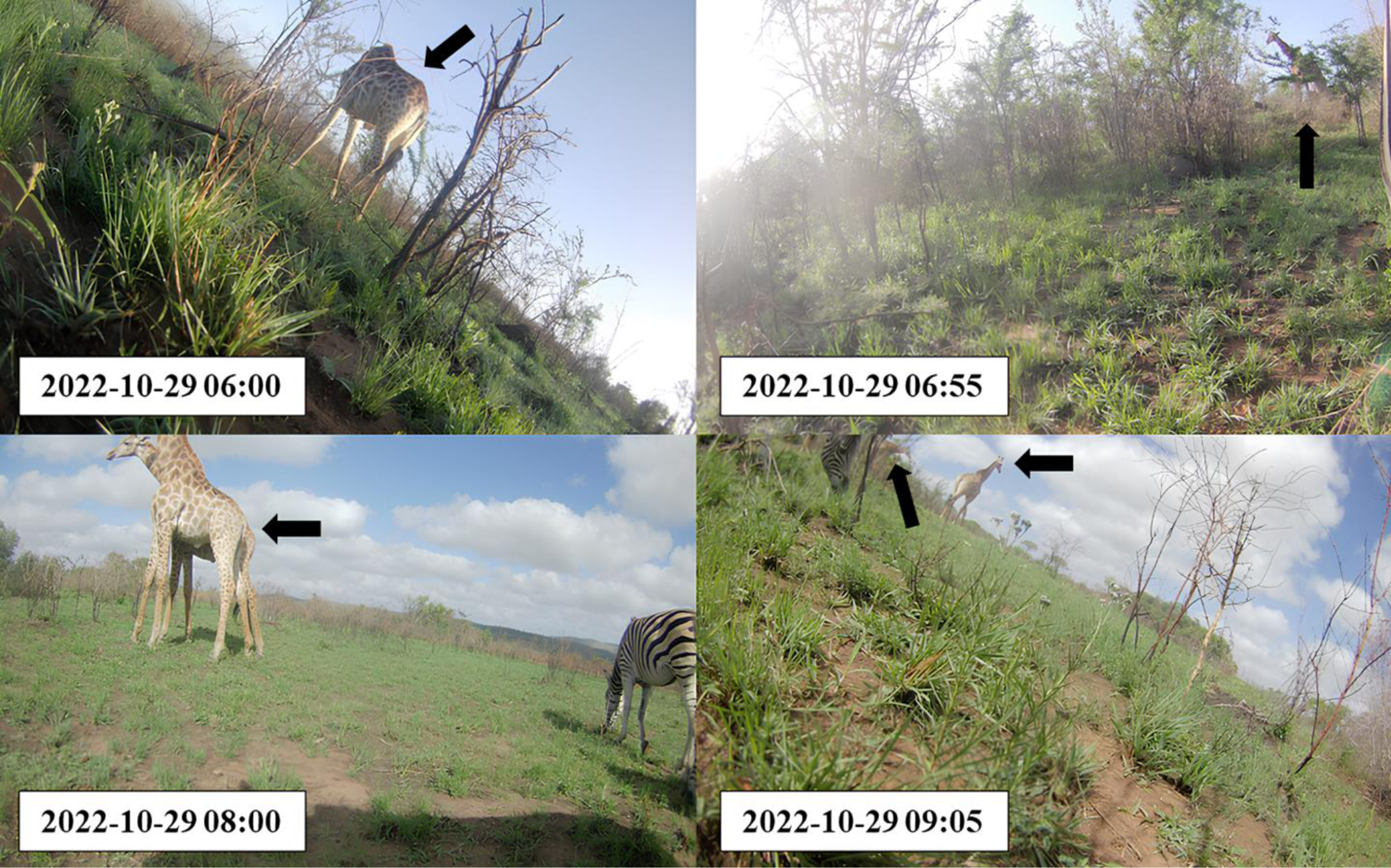
Example of a collared zebra herding with giraffes (pointed by dark arrows) as captured by side-view animal-borne video cameras collecting 10-second videos every 5 minutes.

In savanna ecosystems, herbivores commonly form mixed-species groups to level the landscape of fear (Pays et al. 2014; Beaudrot et al. 2020; Saltz et al. 2023). In particular, zebras are less vigilant when herding with wildebeest, impalas (*Aepyceros melampus*) or giraffes (*Giraffa camelopardalis*) (Schmitt et al. 2014, 2016; Stears et al. 2020). However, little is known about how long they remain together and whether they travel together in the landscape. Here, we describe the fission dynamics of mixed-species herds, and the movement and foraging behaviour of zebras when in such herds. We specifically investigated whether zebras are more encamped when herding with heterospecifics (following the previous findings from Stears et al. (2020) documenting that zebras forage longer time in mixed-species groups) or are more likely to move in mixed-species herds to keep up the rest of a moving group. In the case that zebras travel with heterospecifics, conflicts of interests between moving partners may reduce the cohesion of mixed-species groups. We therefore further explored the link between zebras’ movement decisions and the cohesion of mixed-species groups.

## METHODS

### Study site

The study was conducted in the ∼ 900 km² Hluhluwe-iMfolozi Park, South Africa. In the park, rainfall and vegetation vary greatly with topography, resulting in a gradient between higher elevation (∼750m), higher rainfall (∼975 mm per year) and denser vegetation in the North to lower elevation (∼60m), lower rainfall (∼600 mm per year) and bushed grasslands in the South. The park has a wide diversity of large mammalian herbivores, including giraffes, African buffalo (*Syncerus caffer*), zebras, wildebeest and impalas. A full guild of large mammalian carnivores is also present in the park, including African lions (*Panthera leo*), leopards (*Panthera pardus*), spotted hyaenas (*Crocuta crocuta*), cheetahs (*Acinonyx jubatus*) and wild dogs (*Lycaon pictus*).

### Animal-born video cameras collars

We equipped 6 female zebras, each in a different harem, with side-view camera collars during two capture sessions in October 2022 (n=3) and October 2023 (n=3). Each camera collar recorded 10-second videos every 5 minutes from 05:00 to 18:00 and collected one GPS-location every 15-minute during the day and night time over three weeks. To match each video clip to one zebra location, we interpolated zebra trajectories to 5-minute using a continuous-time correlated random walk model, as implemented in the R package crawl (Johnson et al. 2008). Each of the 17 085 video clips collected in total were observed and manually analysed by R.D. to record whether the tracked zebra was foraging or not and whether it was in a mixed-species group.

### Identification of mixed-species groups

In total, 2 298 out of the 17 085 video clips (13%) showed that the collared zebra was in close proximity (∼50 meters) to heterospecifics. However, animals could be close to the monitored zebra and not be visible on the footage. Indeed, in 6 380 (37%) of the video clips, no other zebra was seen, although we know that individuals of a harem remain together and therefore other zebras were close by (but not seen). We therefore investigated over which interval duration we could assume a species’ presence if not seen (see Appendix S1: Figures S1-S2), and found that we could consider that zebras remained with another species when the time between two video clips showing that species was shorter than one hour. Our results were robust to the use of different temporal thresholds (see Appendix S1: Figure S3 for the sensitivity analyses).

### Study of the fission dynamics of mixed-species groups

To examine the cohesion of mixed-species groups, we conducted a survival analysis of mixed-species groups, modelling the risk of fission of mixed-species groups over time. Since the risk of fission of mixed-species groups may depend on zebras’ movements, we added an interaction term between zebras’ step lengths and whether zebras were with giraffes, impalas, wildebeest or buffalos. We did so by fitting a Cox proportional hazard model using the R package *survival* (Therneau 2015). Since video clips provided information, and hence showed heterospecifics, during daytime only, we controlled for the fact that the observations could be censored, if the monitored zebra was already with heterospecifics one hour after sunrise (15% of mixed-species groups) or if the monitored zebra was still with heterospecifics one hour before sunset (17% of mixed-species groups).

### Study of zebra movements in mixed-species groups

To investigate whether zebras cover long distances when in mixed-species groups, we fitted a Hidden Markov Model (HMM) to step length and turning angles of the movement trajectories to classify each step into a ‘moving’ (large step length and small turning angle) or a ‘encamped’ (small step length and large turning angle) state. We modelled step length using a gamma distribution and turning angle using a Von Mises distribution. To test whether zebras’ movement behaviours could be affected by the presence of heterospecifics, we modelled the transition probability between states as a function of the presence or absence of giraffes, impalas, wildebeest or buffalos. We controlled for diel changes in the probability of being in the ‘moving’ state, independently of heterospecifics, by including hour of the day as a non-linear predictor using splines. Zebra identity was added as a random intercept to account for individual heterogeneity in movement behaviour. This analysis was conducted using the R package *hmmTMB* (Michelot 2023).

### Study of zebra foraging behaviour in mixed-species groups

From each video clip, we classified whether the collared zebra was grazing or not (i.e., standing, scanning the environment, walking), based on the tilted horizon of the image (Figure 1). This approach has been used previously by Kaczensky et al. (2019) to estimate the foraging of khulans (*Equus hemionus*) monitored using video collars. We then investigated whether zebras’ probability of foraging increased when heterospecifics were present, by fitting a mixed logistic regression with the behaviour of the zebra as a binary response (foraging or not) and the presence of giraffes, impalas, wildebeest and buffalos as predictors. Additionally, because zebras’ foraging behaviour in mixed-species groups may depend on their movement behaviours, we added interactions terms between the state identified from the HMM (i.e., encamped or moving state) and the presence / absence of giraffes, impalas, wildebeest or buffalos. As before, we controlled for diel changes in the probability of being foraging, independently of heterospecifics, by including hour of the day as a non-linear predictor using splines. Zebra identity was added as a random intercept to account for individual heterogeneity in foraging rates.

## RESULTS

### General information on mixed-species groups

We observed 373 independent events of zebras in mixed-species groups, mostly with impalas (n = 196) but also with buffalos (n = 80), wildebeest (n = 76) and giraffes (n = 21). On average, collared zebras spent about 27% of their time with another species (16% with impalas, 5% with buffalos, 4% with wildebeest and 3% with giraffes), and less than 2% of their time with more than one species (for example with wildebeest and impalas simultaneously). However, the percentage of time that zebras spent with heterospecifics greatly varied between the monitored harems (5%; 24%; 27%; 28%; 35%; 45%). The lowest value was for a harem in which an adult female gave birth during the tracking period, potentially leading this harem to adopt peculiar spatial and social behaviours. We did not find any clear diel variations in the temporal pattern of zebras’ co-occurrence with impalas, wildebeest, buffalos and giraffes (Appendix S2: Figure S1).

### Fission dynamics of mixed-species groups

On average, zebras remained 1.2 hour with impalas, 0.8 hour with wildebeest, 1 hour with buffalos and 2.2 hours with giraffes. However, the duration of mixed-species groups depended on zebras’ step lengths (Table 1; Appendix S2: Figure S2). Surprisingly, the risks of fission of zebra-wildebeest and zebra-giraffe herds were reduced when zebras moved along longer steps. For a mean step length of zebras, the risk of fission was 0.55 times lower [CI = 0.32; 0.97] when zebras herded with giraffes than with wildebeest and 0.68 times lower [CI = 0.53; 0.86] when zebras herded with impalas than with wildebeest.

**Table 1.** Coefficients (β), hazard ratios (exp[β]) and standard errors (SE) for the Cox model testing the influence of herd mate species and zebras’ step lengths on the risk of fission of mixed-species groups. Positive coefficients show higher risk of fission of the mixed-species groups, relatively to the reference case, here zebra-wildebeest herds. The step length variable is scaled.

| Variables | $\beta$ | $\exp(\beta)$ | SE | z-value | p-value |
| --- | --- | --- | --- | --- | --- |
| **Risk of fission of mixed-species groups** |  |  |  |  |  |
| Zebra – Buffalo | -0,07 | 0,93 | 0,25 | -0,27 | 0,78 |
| Zebra – Impala | -0,39 | 0,68 | 0,12 | -3,21 | < 0.01 |
| Zebra – Giraffe | -0,59 | 0,55 | 0,28 | -2,08 | < 0.05 |
| Step length | -0,15 | 0,86 | 0,06 | -2,52 | < 0.05 |
| Zebra – Buffalo : Step length | 0,18 | 1,20 | 0,08 | 2,23 | < 0.05 |
| Zebra – Impala : Step length | 0,20 | 1,22 | 0,07 | 2,81 | < 0.01 |
| Zebra – Giraffe : Step length | -0,72 | 0,49 | 0,66 | -1,08 | 0,28 |

### Zebra movements in mixed-species groups

The net displacement of zebras was the greatest during periods with giraffes [mean = 300 meters; 90% percentile = 910 meters], lower when with impalas [mean = 240 meters; 90% percentile = 630 meters], and even lower when with buffalos [mean = 140 meters; 90% percentile = 420 meters] or wildebeest [mean = 140 meters; 90% percentile = 310 meters]. Zebras were as likely to shift from moving to encamped states when herding with heterospecifics or when alone (Appendix S2: Table S1; Appendix S2: Figure S3). Similarly, zebras were not more likely to shift from encamped to moving states when herding with wildebeest or buffalos than when with no other species (Appendix S2: Table S1; Appendix S2: Figure S3). This was however observed when they herded with impalas and even more with giraffes, to the point that when with giraffes, zebras were as likely to be moving as encamped (Figure 2), whereas they were more likely to be found encamped in any other situation.

**Figure 2.**
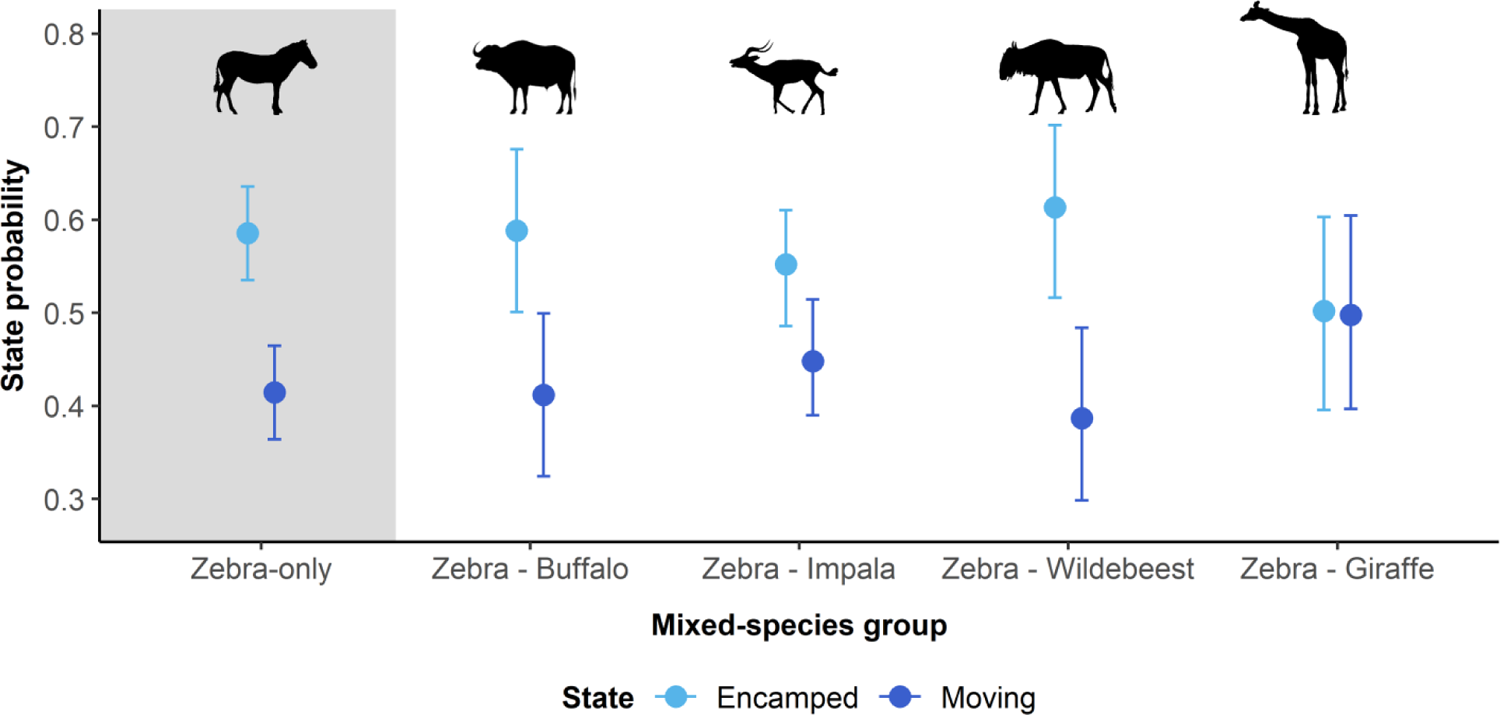
Influence of the presence of other species on the probability that zebras are in the ‘encamped’ or ‘moving’ states. The ‘Zebra-only’ case refers to zebras moving without heterospecifics. Points show the mean parameter estimates and whiskers show 95% confidence intervals.

### Zebra foraging behaviour in mixed-species groups

Being with other species made zebras more likely to forage: they spent 55% [95% CI = 50; 59] of their time grazing when they herded with giraffes, 50% [95% CI = 46; 55] with wildebeest, 49% [95% CI = 45; 53] with buffalos, 49% [95% CI = 46; 52] with impalas and 45% [95% CI = 43; 48] when zebras were without heterospecifics. Additionally, we found that zebras herding without heterospecific, or herding with buffalos, wildebeest or impalas were less likely to forage when moving, but not when they herded with giraffes (Figure 3; Appendix S2: Table S2).

**Figure 3.**
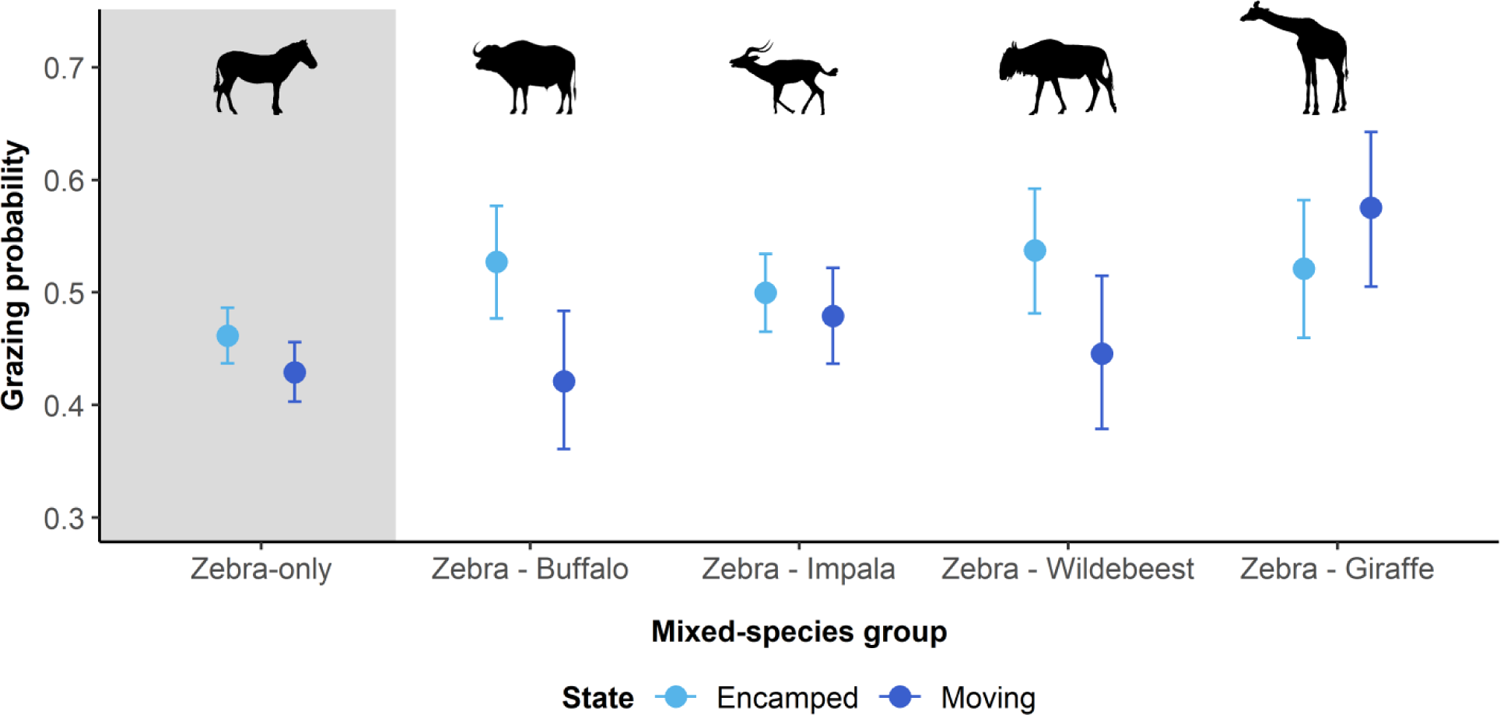
Influence of the presence of other species, and of the movement state of zebras, on the probability to observe zebras grazing. Grazing events were identified based on the tilted horizon of the image recorded by camera collars. The ‘Zebra-only’ case refers to zebras moving without heterospecifics. Points show the mean parameter estimates and whiskers show 95% confidence intervals.

## DISCUSSION

Understanding the intricacies between animal movement and mutualism is a current challenge for ecologists (Shaw et al. 2021). Here, we investigated the movement behaviour of plains zebras in mixed-species groups, showing that they switch more often towards a moving state when herding with giraffes and that this is associated with a lower risk of fission of the mixed-species groups they form with giraffes and a higher grazing probability, compared to when with other species.

Because herding with giraffe is particularly advantageous to decrease the risk of predation, as it allows to eavesdrop on a species that has a great predator detection capacity, (relatively to being without heterospecifics or with impalas, wildebeest or buffalos) (Schmitt et al. 2016), our results suggest that zebras adopt peculiar movement behaviours to ensure the cohesion of zebra-giraffe herds. From a naturalist perspective, Schmitt et al. (2016) anecdotally reported observations of zebras following giraffes to continue to feed in their proximity when they moved off. Although our results suggest that zebras adjust their movement behaviours to follow giraffe and forage in their proximity, herding events with giraffes were still rare and mixed-species grouping was more often observed with impalas rather than giraffes. Because impalas are about 10 times more abundant than giraffes in Hluhluwe-iMfolozi Park (le Roux et al. 2017), such differences in the number of herding events likely reflect differences of herd mate availability. Additionally, the low abundance of giraffes may strengthen the adjustments of zebras’ movement behaviours: zebras may adopt peculiar movement pattern to stay as long as possible in proximity to a rare partner that has a great predator detection capacity.

Following giraffes may ensure higher abilities to detect predators. However, such behaviour could be energetically costly if it reduces the time available for zebras to forage. Here, on the contrary, our results show that zebras spent longer time grazing in mixed-species groups, especially when herding with giraffes, supporting the previous findings from Stears et al. (2020). Surprisingly, the increased time devoted to forage when herding with giraffes was not restricted to encamped but also to moving states, whereas increased foraging time when herding with impalas, wildebeest or buffalos were mostly correlated to encamped movements of zebras.

Because species may have different energetic needs, may select habitat differently, or may respond differently to the risk of predation, maintaining group cohesion is challenging and individuals should try to maintain cohesion on the long-run only if the benefits of doing so are high. Here, we found that the cohesion of mixed-species group was lower when zebras herded with wildebeest or buffalos than with impalas or giraffes. Conflicts of interests in decision making lower the cohesion of single-species group, such as observed in red deer (*Cervus elaphus*) or woodland caribou (*Rangifer tarandus*) (Conradt and Roper 2000; Sueur et al. 2011; Le Goff et al. 2024), and can be expected to constrain even more strongly the cohesion of mixed-species group. Although zebras benefit from a lower risk of predation by herding with wildebeest than impalas (Schmitt et al. 2014), the lower difference of habitat selectivity with impalas than wildebeest may explain the higher cohesion of zebras herding with impalas. Wildebeest are indeed more selective grazers than zebras, and their movement decisions may be differently constrained by energetics needs, potentially leading to conflicts of interests that increase the risk of fission.

## CONCLUSION

Here, we investigated the movement of plains zebras in mixed-species groups, providing new insights on prey mutualism in savanna ecosystems. Our results show that using animal-borne video collars is an adequate tool to explore the movement of mixed-species groups, offering promising perspectives on the intricacies between animal movement and positive interspecific interactions. An important challenge for ecologists is to study collective decision making in mixed-species groups (Sridhar and Guttal 2018; Oliveira and Bshary 2021). However, because animal-borne video collars allow us to detect the presence of heterospecifics but not their movements, further questions relative to collective decision making remain unfortunately unanswered, such as which species departs first or which species is the leader or the follower.

## ACKNOWLEDGEMENTS

We are grateful to Ezemvelo for providing the authorization to conduct this research, and particularly to D.J. Druce and S. Mbongwa for their continuous administrative and field support. We acknowledge the field efforts of B. Gehr, Y. Richard, R. Leeming, J. Lawrence, M. Krings and anyone involved in the zebra captures. This study was partly funded by the TERRA FORMA project (ANR-21-ESRE-0014), the REPOS project funded by the i-Site MUSE (ANR-16-IDEX-0006), the SOCIALIPOP project (ANR-22-CE02-0011-02) and the FutureFear project sponsored by the Fondation BNP Paribas.

## AUTHOR CONTRIBUTIONS

SCJ initiated the zebra research project, acquired funding and coordinated data acquisition. SCJ and RD conceived the ideas and designed the statistical methodology. RD collected data and conducted the statistical analyses. RD, SCJ and MV interpreted the results. RD led the writing of the manuscript. RD, SCJ and MV revised and edited the manuscript.

## Appendix S1. Finding the interval over which assuming a species’ presence if not seen is correct

Because animals could be close to the monitored zebra and not be visible on the footage (side-view video), we needed an approach to define whether zebras were still in mixed-species herds when we recorded the presence of another species but that species has not been seen on video clips for a certain time. To do so, we decided to use a temporal threshold below which we could assume a species’ presence if not seen (under the condition that the species was seen a certain time ago). To use a relevant temporal threshold, we used three approaches:

First, R.D. conducted field observations to record whether other species were around the collared zebra. Observations lasted as long as collared zebras were visible in the field. When zebras were known (from field observations) to herd with heterospecifics over a certain period (for example herding with wildebeest during 2h), we compared how much video clips were needed to detect the presence of wildebeest around the collared zebras (from collar observations) (Figure S1). Doing so, we found that after 1h of video clip observations, at least one video clip showed the heterospecific i.e., inversely, after 1h of video clip observations, if no videos showed heterospecific it is likely that no other species was around the collared zebra.

Second, we used several temporal thresholds (from 5 minutes to 3 hours) to assume a species’ presence over a certain time after the last detection of this species. Because 6 380 (37%) of the video clips did not show other zebras, and because individuals of a harem remain together all day long, we used these temporal thresholds to infer how many independent encounters with other zebras we could detect per day (i.e., encounters spaced by a longer time period than the temporal threshold), and how long this encounter lasted. We further expect to use a temporal threshold allowing us to detect one encounter event per day with zebras lasting ∼13 hours. On the contrary, detecting more than one encounter with zebras would indicate that the corresponding temporal threshold does not allow us to interpolate the presence of zebras not seen on video clips (and that are known to be in the harem of the collared zebra). Figure S2 shows that using a temporal threshold higher than 1 hour is needed; otherwise, we frequently consider temporal periods when zebras are assumed to move alone, whereas other zebras are still around them.

Third, we used the different temporal thresholds (from 5 minutes to 3 hours) to compare the number of independent encounters with heterospecifics identified from video clips. Figure S3 shows that using a temporal threshold higher than 1 hour is needed; otherwise, the number of encounters is not at a plateau: encounters with heterospecifics are not independent.

**Appendix S1: Figure S1.**
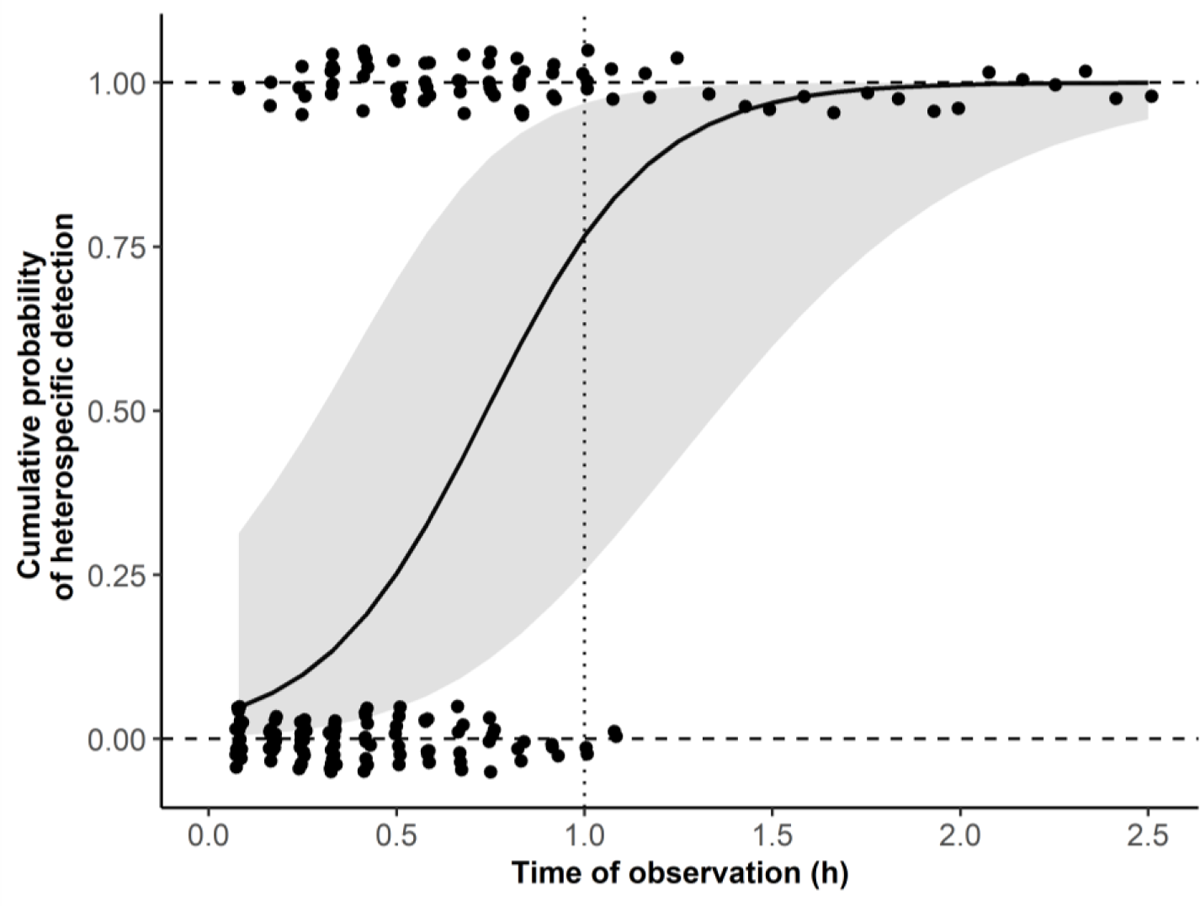
Cumulative probability to detect the presence of a mixed-species herd from video clips when the presence of heterospecific was known from field observation simultaneously to collar deployment.

**Appendix S1: Figure S2.**
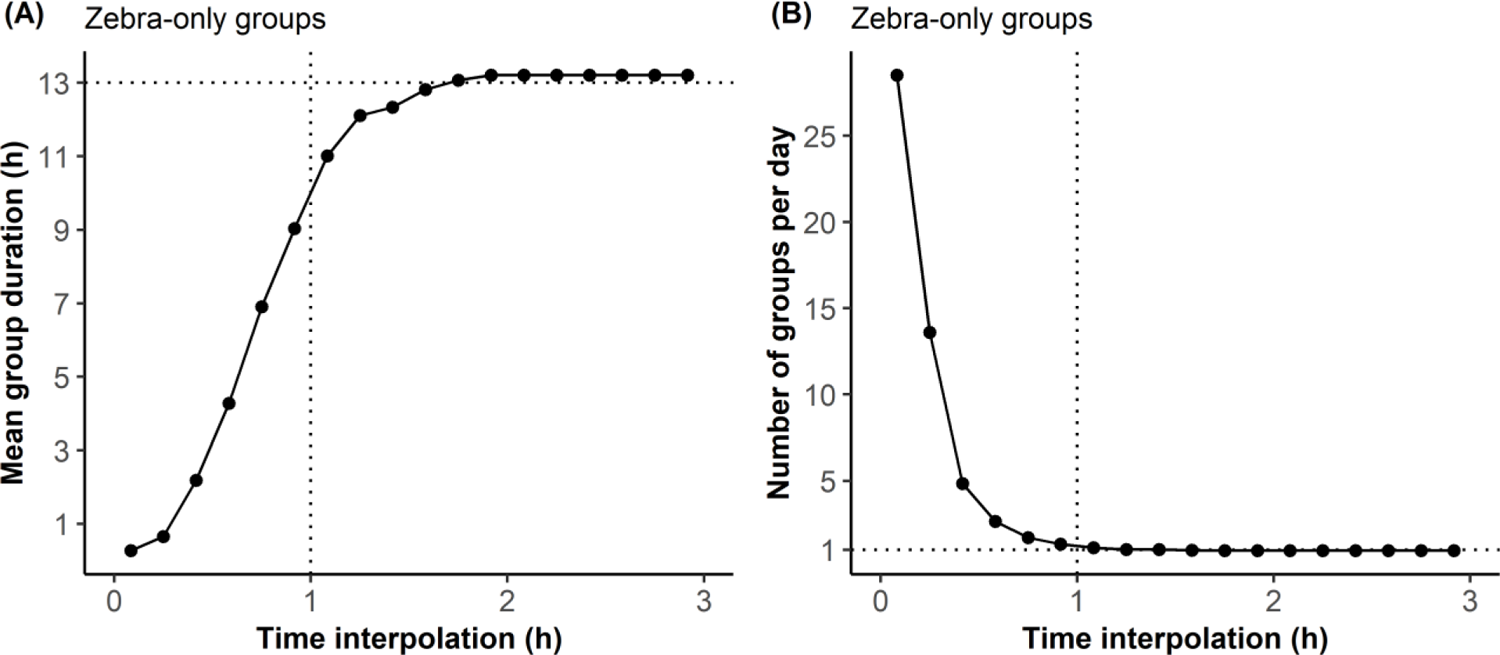
(A) Mean duration and (B) number of zebra-only groups detected from video clips according to the time of interpolation used to assess that a zebra is still present around the collared zebra even when not visible on video clips. Because zebra harems are cohesive, we expect to observe only one zebra-only herd event per day, throughout the day (∼13 hours).

**Appendix S1: Figure S3.**
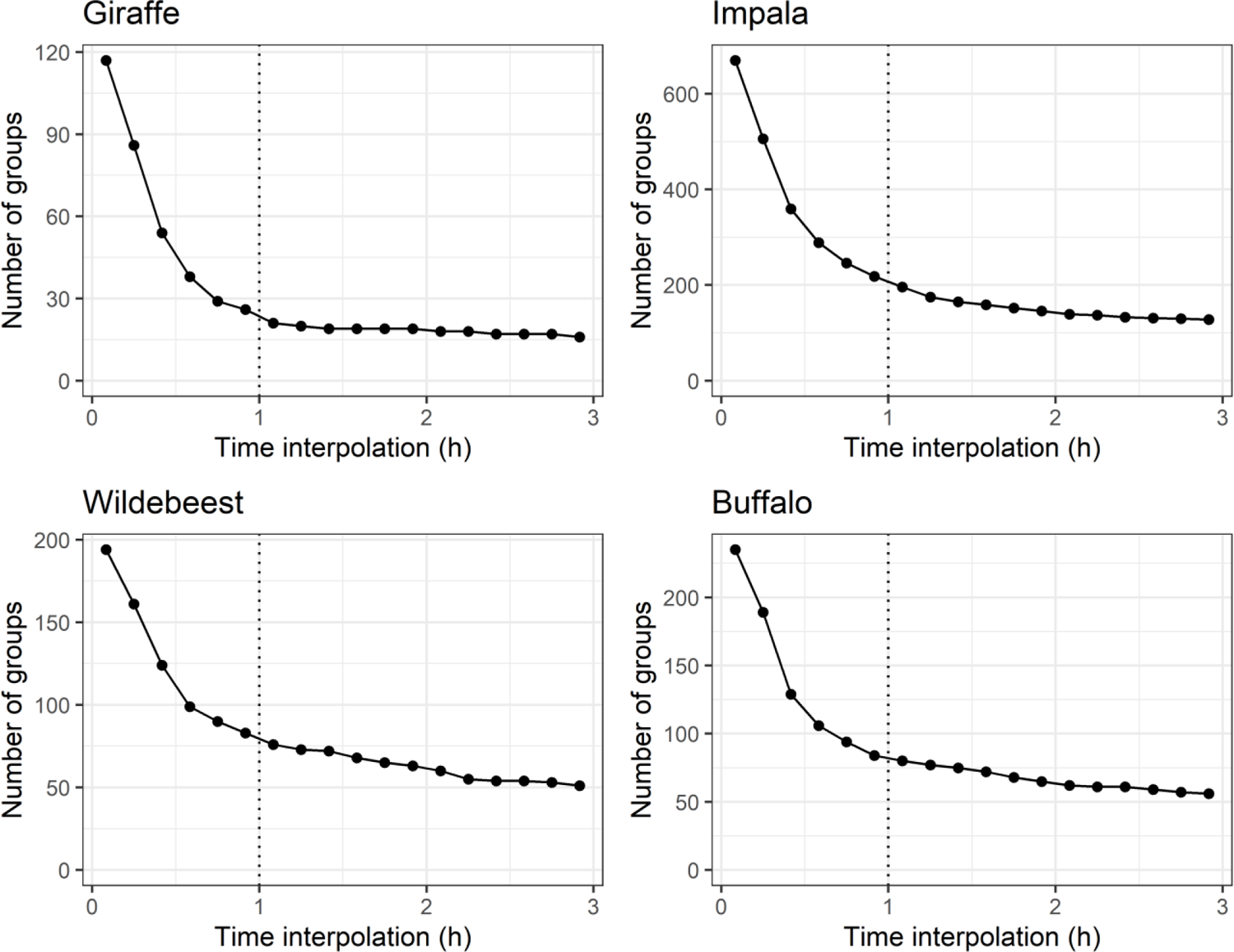
Number of zebra-wildebeest herds, zebra-giraffe herds, zebra-impala herds and zebra-buffalo herds according to the time of interpolation used to assess that a species is still present around the collared zebra even when not visible on video clips.

**Table S1.**
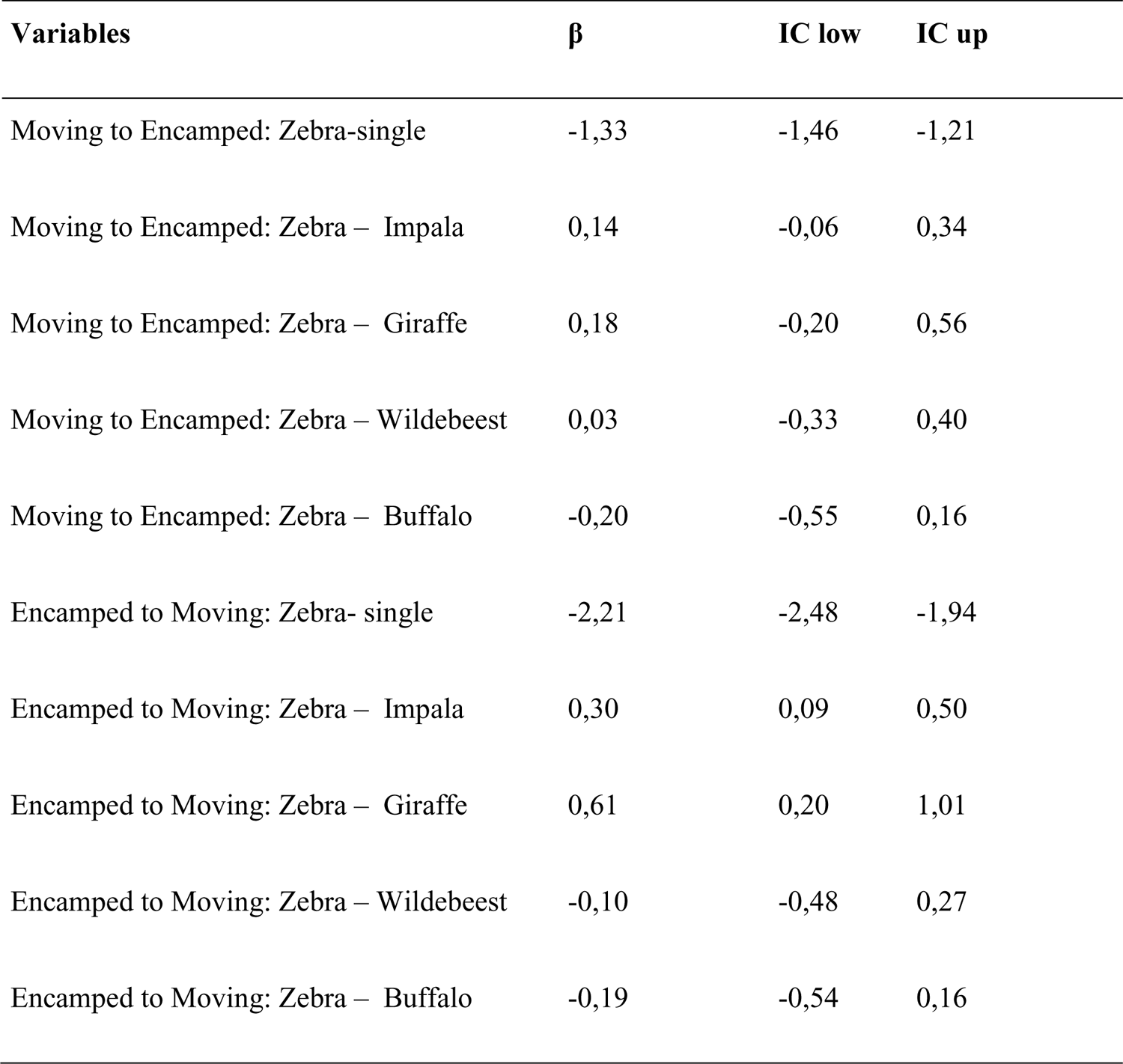
Statistics of the model testing the influence of being in a mixed-species herd on the probability that zebras shifted from moving to encamped or, conversely, from encamped to moving states.

| **Variables** | **$\beta$** | **IC low** | **IC up** |
| --- | --- | --- | --- |
| Moving to Encamped: Zebra-single | -1,33 | -1,46 | -1,21 |
| Moving to Encamped: Zebra – Impala | 0,14 | -0,06 | 0,34 |
| Moving to Encamped: Zebra – Giraffe | 0,18 | -0,20 | 0,56 |
| Moving to Encamped: Zebra – Wildebeest | 0,03 | -0,33 | 0,40 |
| Moving to Encamped: Zebra – Buffalo | -0,20 | -0,55 | 0,16 |
| Encamped to Moving: Zebra- single | -2,21 | -2,48 | -1,94 |
| Encamped to Moving: Zebra – Impala | 0,30 | 0,09 | 0,50 |
| Encamped to Moving: Zebra – Giraffe | 0,61 | 0,20 | 1,01 |
| Encamped to Moving: Zebra – Wildebeest | -0,10 | -0,48 | 0,27 |
| Encamped to Moving: Zebra – Buffalo | -0,19 | -0,54 | 0,16 |

**Table S2.** Statistics of the model testing the influence of being in a mixed-species herd and of the movement state of zebras on the probability to observe zebras grazing.

| **Variables** | **$\beta$** | **SE** | **z-value** | **p-value** |
| --- | --- | --- | --- | --- |
| Zebra-single | 0,47 | 0,17 | 2,76 | < 0.01 |
| Zebra – Impala | 0,15 | 0,06 | 2,63 | < 0.01 |
| Zebra – Giraffe | 0,24 | 0,12 | 1,92 | 0.06 |
| Zebra – Wildebeest | 0,30 | 0,11 | 2,81 | < 0.01 |
| Zebra – Buffalo | 0,26 | 0,09 | 2,81 | < 0.01 |
| Moving | -0,13 | 0,04 | -3,24 | < 0.01 |
| Zebra – Impala: Moving | 0,05 | 0,10 | 0,50 | 0.62 |
| Zebra – Giraffe: Moving | 0,35 | 0,19 | 1,84 | 0.07 |
| Zebra – Wildebeest : Moving | -0,24 | 0,17 | -1,36 | 0.17 |
| Zebra – Buffalo: Moving | -0,30 | 0,15 | -1,95 | 0.05 |

## Appendix S2. Additional tables and figures

**Appendix S2: Figure S1.**
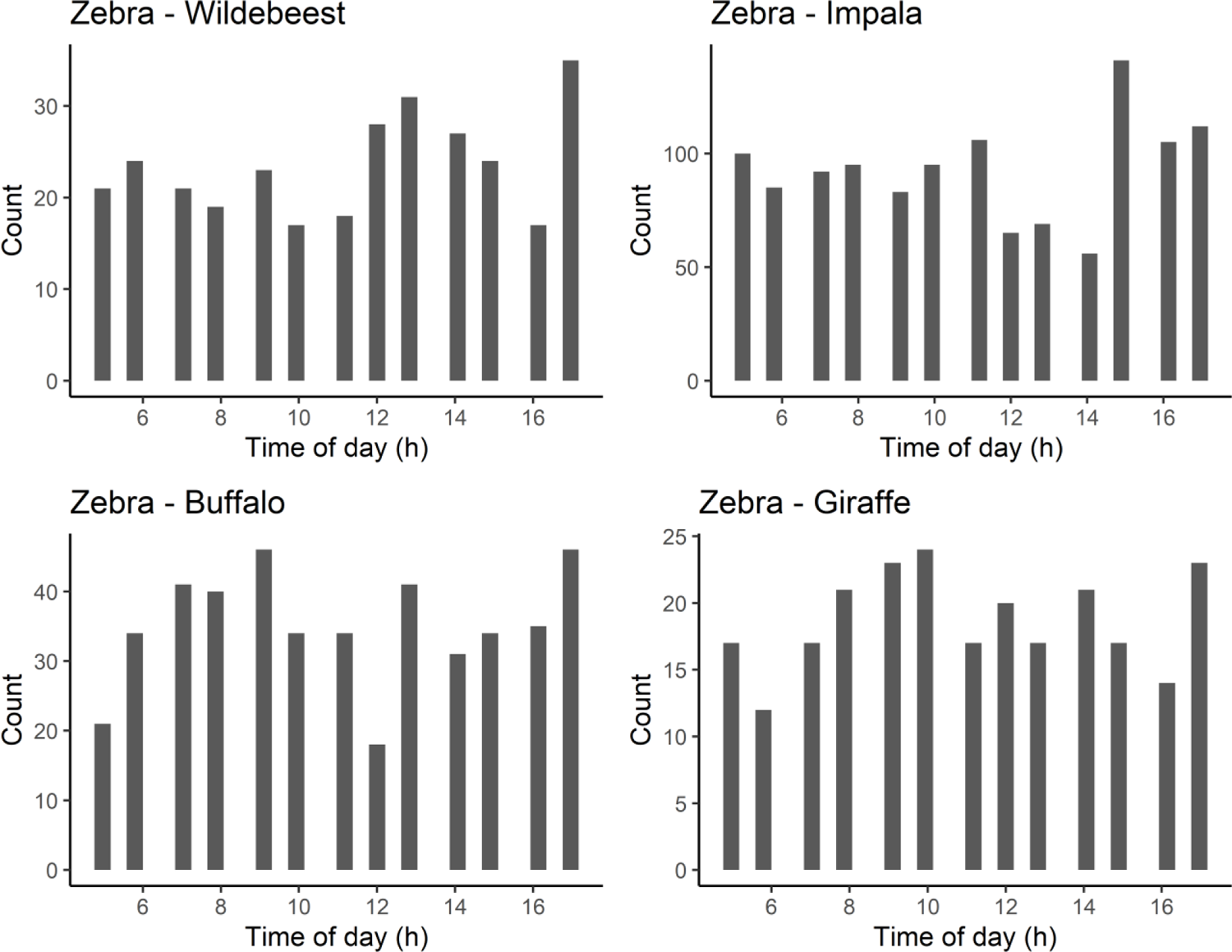
Distribution of mixed-species herds observations over the day. Each bar shows the number of video clips showing the corresponding species according to the hour of the day. In total, we obtained 315 video clips in which wildebeest could be seen, 1 264 video clips with impalas, 472 video clips with buffalos and 247 video clips with giraffes.

**Appendix S2: Figure S2.**
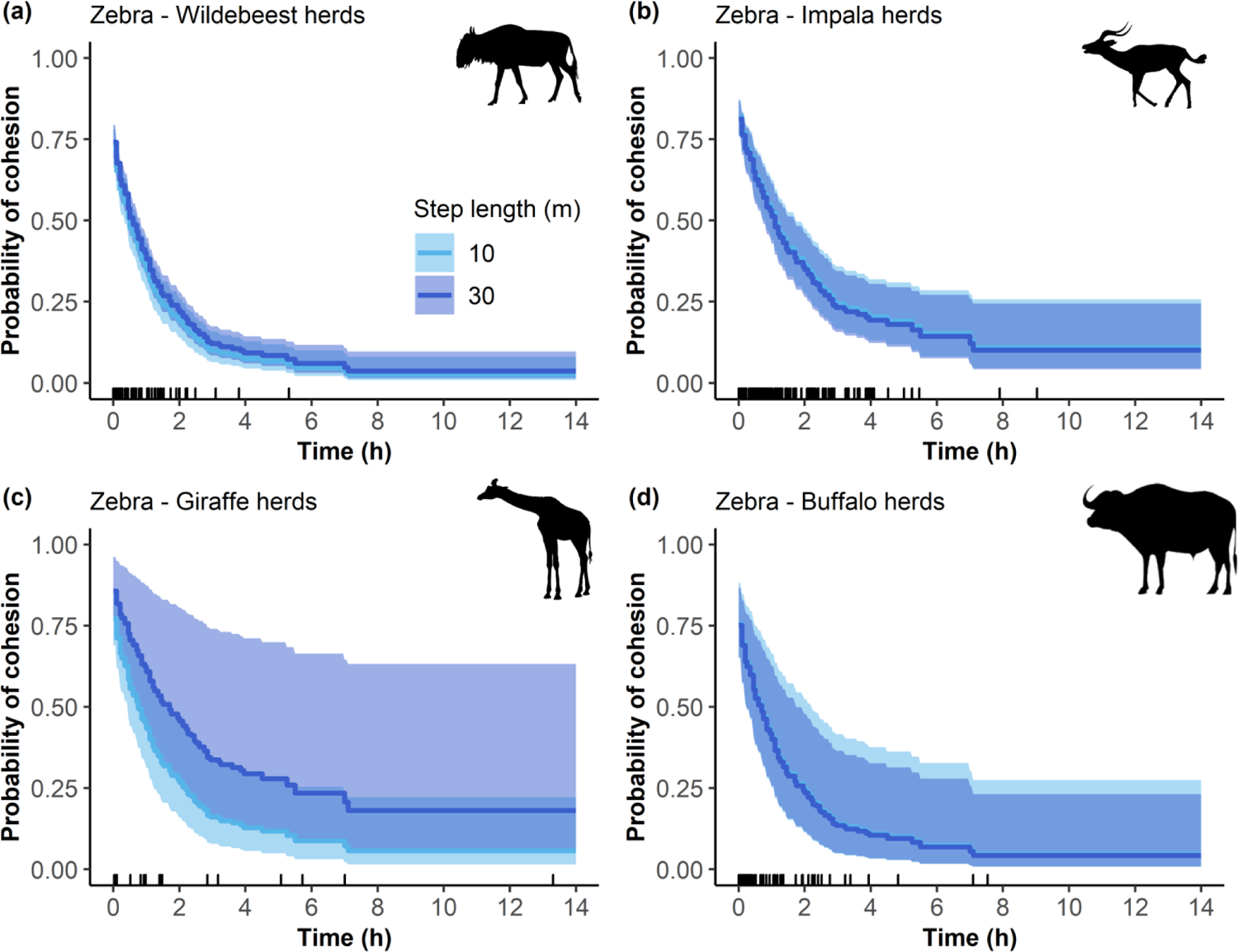
Temporal changes in the probability of cohesion of (a) zebra-wildebeest herds (b) zebra-impala herds (c) zebra-giraffe herds (d) zebra-buffalo herds. Fission events of zebra-giraffe herds were less likely when zebras moved along long distances (dark blue) than short distances (light blue) (p-value < 0.001; Table S). Time start at 0 is the onset of fusion of mixed-species herds. Ribbon extremities show 95% confidence interval, whereas lines show the mean probability of cohesion of mixed-species herds. Small vertical lines show actual fission events observed.

**Appendix S2: Figure S3.**
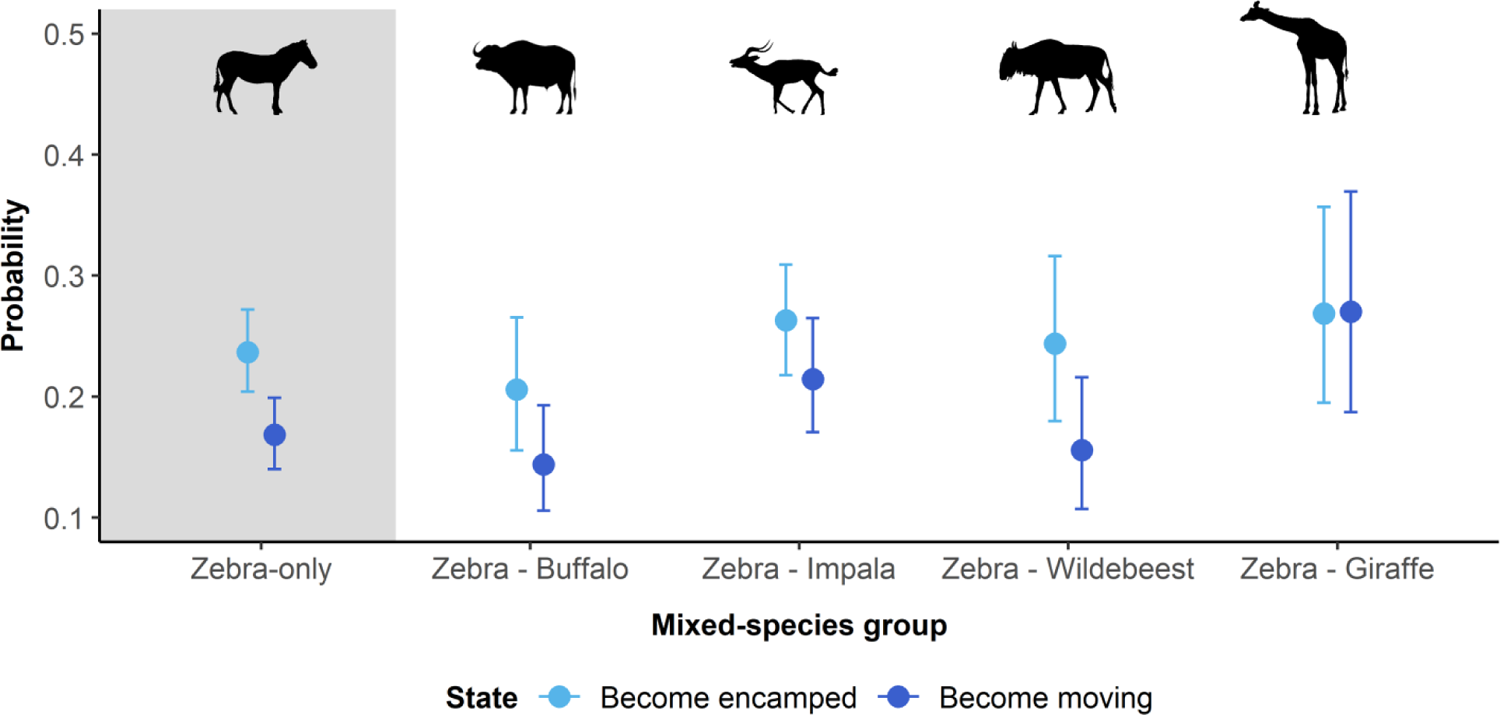
Transition probability (1) from the moving to the encamped state and (2) from the encamped to the moving state as a function of the species zebras herd with. The ‘Zebra-only’ case refers to zebras moving without heterospecifics. Points show the mean parameter estimate and whiskers show 95% confidence intervals.

